# Multiquantal transmission supports dynamic synaptic strength in vivo

**DOI:** 10.64898/2026.10.18.759916

**Authors:** Paras A. Patel, Michael J. Higley

## Abstract

In the central nervous system, synaptic transmission occurs through the probabilistic release of neurotransmitter vesicles following presynaptic action potentials (AP). Traditional models posit a single vesicle release event per AP, though experimental evidence indicates multivesicular release (MVR) may occur under some circumstances, greatly expanding the dynamic range and information content at single synapses. However, the quantal nature of synaptic transmission in the intact, waking brain is largely unknown. Here, we used two-photon imaging of iGluSnFR4f-expressing cortical axons and found that MVR dominates synaptic transmission at both layer 4 and layer 2/3 synapses, enhancing reliability and yielding non-binary signaling for single synaptic connections. Moreover, MVR in the presence of cholinergic modulation conveys behavioral state-dependent influence over synaptic potency. Overall, our findings demonstrate that MVR at central synapses in vivo contributes to the computational capacity of cortical circuits.

## Introduction

Synaptic transmission forms the basis of communication between neurons in the nervous system. In the canonical model, axonal action potentials invade the presynaptic terminal, inducing the voltage- and calcium-dependent fusion of synaptic vesicles that release their contents into the synaptic cleft for sensing by the postsynaptic cell (*1, 2*). However, vesicle fusion is inherently probabilistic and is known to vary across cell types and in response to activity and neuromodulatory influences (*3, 4*). Early studies suggested that for each presynaptic action potential, a single vesicle could be released (*5, 6*). In this case, transmission is binary (failure or success of release) with synaptic potency (the magnitude of the postsynaptic response given a successful release) determined primarily by postsynaptic mechanisms like receptor density (*7*). However, several more recent studies found that under some conditions, central synapses in the neocortex, hippocampus, and cerebellum could exhibit multivesicular release, where a single action potential triggers synchronous fusion of more than one vesicle (*8*–*13*). In this case, synaptic potency is also influenced by the number of vesicles at the terminal, with additional vesicles producing a larger response assuming lack of receptor saturation (*13*–*15*).

In the neocortex, glutamatergic synaptic transmission forms the fundamental mechanism by which cortical circuits respond to, amplify, and filter signals underlying perception, cognition, and behavior (*16*–*18*). Surprisingly, the nature of transmission at single synapses, particularly in vivo, is poorly understood. Ex vivo studies have generally found that cortical synapses exhibit low reliability stemming from low release probability (*19*–*22*), shaping models of network function (*23*–*25*). However, experimental challenges have largely prevented quantitative assays of single synapse function in the cortex of awake animals, limiting a more thorough understanding of synaptic function.

## Results

### Glutamate release is multiquantal in vivo

To monitor synaptic release at single glutamatergic boutons in vivo, we used a viral vector to drive sparse expression of the fluorescent glutamate sensor iGluSnFR4f in pyramidal neurons (PNs) of mouse visual cortex. Layer 2/3 PNs were labeled by co-injection of AAV9-FLEX-CAG-iGluSnFR4f and AAV9-CaMKII-Cre (see Methods). In separate experiments, we labeled layer 4 PNs using viral expression of Cre-dependent iGluSnFR4f in transgenic *Scnn1a*-Cre mice (*26*). Injected mice were then imaged during waking head-fixation using 2-photon microscopy at high frame rates (250-280Hz) (Figure 1A). Axons from L4 PNs were identified as smooth, narrow caliber fibers located in more superficial areas above their corresponding somatodendritic domains. L2/3 axons were interspersed with labeled dendrites but were easily distinguished by the lack of spines. For both types of cells, high magnification imaging revealed varicosities confirmed to be active synapses by the presence of spontaneous, spatially restricted fluorescence transients (Figure 1B-C).

**Figure 1:**
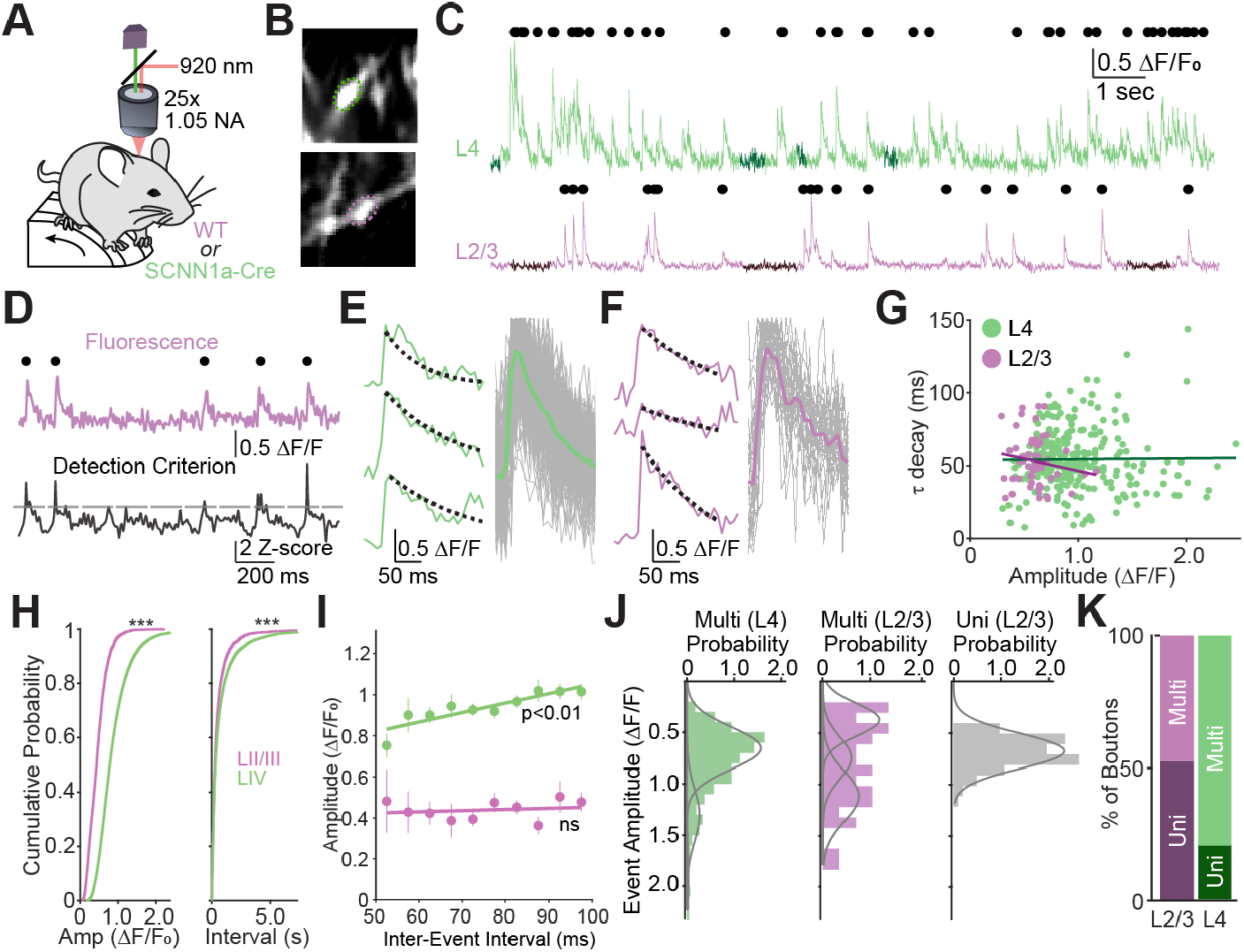
(A) Schematic of experimental setup illustrating either WT C57BL/6J or Scnn1a-Cre transgenic animals free to run on a wheel while head-fixed and two photon excitation/emission paths. (B) Example of two individual boutons imaged either from layer IV axons (top) or layer II/III axons (bottom). (C) Traces from the boutons in (B) with detected events labeled as circles. Dark regions of the trace indicate event-free epochs used to calculate noise. (D) Example of event detection indicating raw fluorescent trace (top) and calculated detection criterion aligned to the trace in time (bottom). (E) example of three individual events from a layer IV bouton with single exponential decay fit from the peak onwards overlaid (left) with all events from that bouton overlaid and the mean waveform indicated as a colored trace (right). (F) Same as (E) but for a layer II/III bouton. (G) Tau decay values for each event from the boutons in (E-F) plotted against amplitude with a linear fit showing no significant relationship for either bouton (L23 r=-0.15, p=0.25; L4 r=0.01, p=0.86). (H) Amplitude histograms across all imaged boutons for both cell populations (n=48 boutons/3 animals layer 2/3, mean 0.46±0.03 ΔF/F_0_; n=109 boutons/6 animals L4, mean 0.83±0.03 ΔF/F_0_, p=5.75×10^-14^). The inter-event interval distributions for the same boutons is plotted on the right (layer 2/3 mean 0.69±0.06s, layer 4 mean 1.13±0.08s, p=1.1×10^-4^). (I) Mean amplitude at 5ms interval bins at short intervals. Layer 4 Spearman’s correlation r=0.89, p=0.00138; layer 2/3 r=0.16, p=0.657. (J) Example amplitude histograms from single layer IV and layer II/III boutons classified as multiquantal (left, middle) and an example uniquantal bouton from layer II/III (right). (K) Proportion of boutons classified as uniquantal or multiquantal (47% L23 classified multiquantal, 79% L4 classified multiquantal).

Surprisingly, the amplitudes of these spontaneous transients varied several-fold at individual sites, suggesting that glutamate release did not follow an all-or-none distribution (Figure 1C). To quantify these signals, release events were isolated using a template-based detection algorithm (Figure 1D, see Methods). The decay kinetics for individual events was highly consistent, matching *ex vivo* values (*27*), and did not correlate with event magnitude (Figure 1E-G), suggesting they arose from highly synchronous release at single sites. Variable event amplitude might also arise from pooling of glutamate from neighboring release sites mistakenly labeled as one. However, our imaging point spread function is ∼0.5 micrometers in the x-y plane, and data from the publicly available MICrONS electron microscopy dataset (*28*) indicate that <20% of synaptic boutons are within 1 micrometer of their nearest neighbor, arguing against this possibility.

We compared the distribution of amplitudes between the two cell types, finding that L4 PNs exhibited significantly larger events versus L2/3 (mean layer 4 amplitude 0.83±0.03, mean layer 2/3 amplitude 0.46±0.03, p=5.75×10^-14^ two-sample ranksum on per-bouton means). The frequency of L4 events was also significantly lower (mean layer 4 interval 1.12±0.08s, mean layer 2/3 interval 0.69±0.06s, p=1.1×10^-4^ two-sample ranksum on per-bouton means)(Figure 1H). The amplitude of L4 events was also significantly correlated with the inter-event interval (Spearman r=0.89, p=0.0014), suggesting that recent synaptic history can influence the amount of glutamate released (Figure 1I). Interestingly, L2/3 events did not show a similar time-varying change in amplitude (r=0.16, p=0.657, Figure 1I).

We next plotted the distribution of event amplitudes from single boutons and iteratively fit a varying number of gaussian curves to each histogram (Figure 1J). Boutons were classified as multiquantal if the distribution was significantly better fit with multiple versus single gaussians by both Likelihood Ratio Test and Bayesian Information Criterion. With this approach, we observed both multiquantal and uniquantal boutons from our data set. Moreover, layer 4 PNs exhibited a significantly greater fraction of multiquantal boutons versus layer 2/3 PNs when measured from spontaneous data (79% vs. 47%, p=6.8×10^-4^ Fisher’s Exact Test)(Figure 1K). Overall, our data indicate that glutamatergic transmission in vivo exhibits substantial event-to-event variation in the amount of glutamate release at single synapses, consistent with the presence of multivesicular release for both L4 and L2/3 axons.

### Quantal analysis reveals multivesicular release for high- and low-Pv synapses

Spontaneous synaptic events preclude detailed quantal analysis as the underlying presynaptic spike train is unknown. Therefore, we modified our experimental protocol to include a chronically implanted stimulating electrode near the imaging field of view (Figure 2A). Separate experiments in which we imaged axonal calcium transients revealed that single axons were extremely reliable in response to trains of electrical stimulation (10 Hz), showing essentially no failures (data not shown). We then paired stimulation with iGluSnFR4f imaging at single L4 or L2/3 boutons, finding that synaptic events could also be elicited in response to stimulation (Figure 2B-C). However, a subset of stimuli failed to evoke a glutamate transient, indicating failure to release on those occasions (Figure 2C), consistent with the probabilistic nature of synaptic transmission.

**Figure 2.**
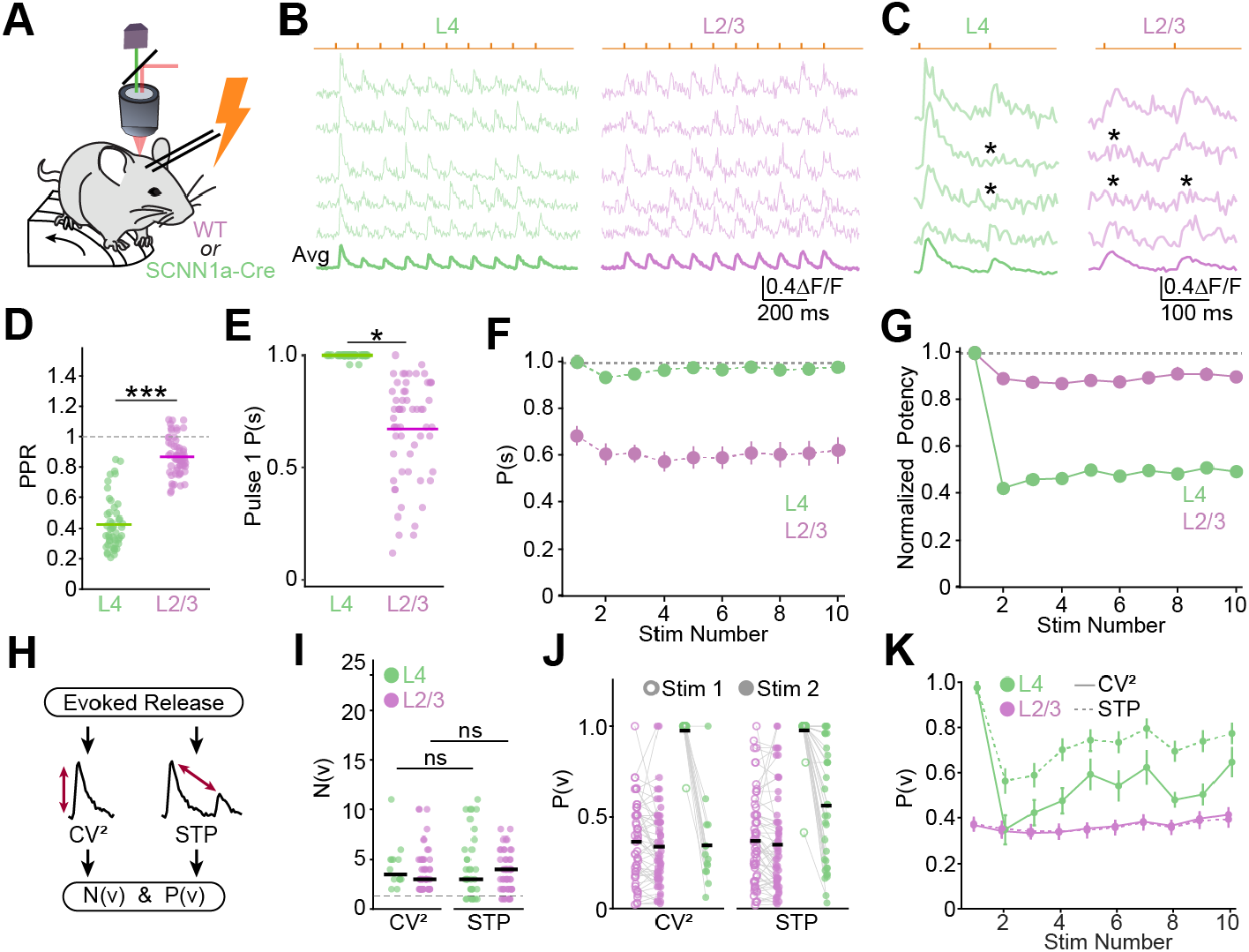
(A) Schematic of experimental diagram illustrating *in vivo* local electrical stimulation. (B) Example trials from layer IV (left, green) and layer II/III (right, pink) boutons demonstrating amplitude variation in single trials with the average across 25 trials below. Stimulus times are indicated by the orange tick marks. (C) Isolated pulse 1 and 2 single trials from the boutons in (B) with average below. Asterisks indicate release failures. (D) Paired pulse ratio including failed trials between pulse 1 and 2 for each population (L23 mean 0.8672 +/- 0.0156, L4 mean 0.4263 +/- 0.0241; p = 4.18×10^-17^ two-sample ranksum test). (E) Average first pulse success rate per bouton (L23 mean 0.6675 +/- 0.028 (n=61 boutons/5 animals), L4 mean 0.9984 +/- 0.00114 (n=49 boutons/5 animals); p = 2.90×10^-20^ two-sample ranksum test). (F) Average success rate across all 10 stimulation pulses in the train. (G) Normalized amplitude excluding failures across the 10Hz train for each population (potency). (H) Schematic of independent strategies for N(v) and P(v) estimation. (I) Calculated N(v) values across methods and cell populations (L23 CV2 mean 3.8±0.3 (n=56 boutons/4 animals), L23 STP mean 3.6±0.2 (n=61 boutons/4 animals), p=0.861 two-sample ranksum test, L4 CV2 mean 4.3±0.6 (n=13 boutons/2 animals), L4 STP mean 4.3±0.5 (n=30 boutons/2 animals), p=0.543 two-sample ranksum test). (J) Calculated P(v) across methods and populations (L23 CV2 pulse 1 mean 0.36±0.03, L23 CV2 pulse 2 mean 0.33±0.03, L23 STP pulse 1 mean 0.37±0.03, L23 STP pulse 2 mean 0.35±0.03, L4 CV2 pulse 1 mean 0.98±0.02, L4 CV2 pulse 2 mean 0.35±0.06, L4 STP pulse 1 mean 0.98±0.02, L4 STP pulse 2 mean 0.57±0.06). (K) Calculated P(v) across all 10 pulses for each bouton population and method.

We first calculated the paired pulse ratio, a standard measure of short-term plasticity, as the ratio of average response amplitudes for the second versus first stimulus in the train. L4 boutons exhibited significant paired-pulse depression at a 100ms interval (mean PPR 0.43±0.02), while L2/3 boutons exhibited only modest depression (mean PPR 0.87±0.02; p=4.18×10^-17^ two-sample ranksum test) (Figure 2D). For univesicular models of release, paired-pulse depression reflects a decrease in the fraction of successful release trials. However, our analyses revealed the surprising finding that neither L4 nor L2/3 boutons exhibited a change in the average success rate between pulses (Figure 2E-F). Instead, we observed a reduction in synaptic potency (average amplitude for successful release trials) for both L4 and L2/3 (Figure 2G). These results are instead consistent with multivesicular release, where short-term changes in *Pv* correspond to a reduction in the number of vesicles released.

For a bouton with *N* vesicles that fuse independently with a given probability *Pv* per each action potential, the number of fused vesicles per event follows a binomial distribution (*29*–*32*). In this case, the coefficient of variation of event amplitudes is proportional to 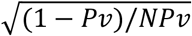. In addition, the probability that a vesicle fails to fuse on a given trial, *P*_*F*_ = (1 − *Pv*)^*N*^. Together, these equations allow us to estimate the parameters *N* and *Pv* for each bouton (Figure 2H). With this approach, the average *N* was similar between layers with a mean of 4.3±0.6, range 1-11, for L4 and 3.8±0.3, range 1-10, for L2/3 boutons (p=0.17 two-sample ranksum test). Furthermore, the mean calculated *Pv* for the first stimulus in a train was 0.98±0.02 for L4 boutons and 0.36±0.03 for L2/3 boutons (p=1.85×10^-8^, two-sample ranksum test) (Figure 2I-J).

Our experimental protocol also allows us to compare transients over the course of the stimulus train, where we assume the number of total potential vesicles *N* does not change. In this case, the change in average amplitudes must reflect variation in *Pv*. This relationship can also be combined with synaptic failure rate to yield an independent estimate of *N* and *Pv*, producing values that did not differ significantly from those obtained using the coefficient of variation (Figure 2I-K). Overall, our data demonstrate that layer 2/3 boutons exhibit a low P(v) but minimal use-dependent change in synaptic potency. In contrast, layer 4 boutons exhibit extremely high P(v), marked reduction in synaptic potency for individual events, consistent with reduction in vesicles released rather than failure of synaptic transmission. Overall, these analyses indicate that multivesicular release can be observed at cortical synapses with either high or low basal *Pv*.

### Quantal variation for synapses along the same axon

Synaptic quantal parameters might be similar for a given presynaptic cell, suggesting that a neuron coordinates signals across its outputs. Conversely, individual synapses might be tuned independently to maximize computational flexibility. We compared these two models by examining a subset of recordings in which we obtained multiple boutons along the same axon (Figure 3A). Results showed that for both L4 and L2/3 PNs, values for *N* and *Pv* could vary considerably for nearby boutons (Figure 3A). We therefore calculated the difference in *N* or *Pv* for all possible pairs of same-axon boutons (Figure 3B). Surprisingly, we found no significant difference between *ΔN* and *ΔPv* measured for same-axon pairs compared to either different-axon pairs or a shuffled control (Figure 3B). Thus, our results suggest an independent tuning model for presynaptic quantal variation within the cortex.

**Figure 3.**
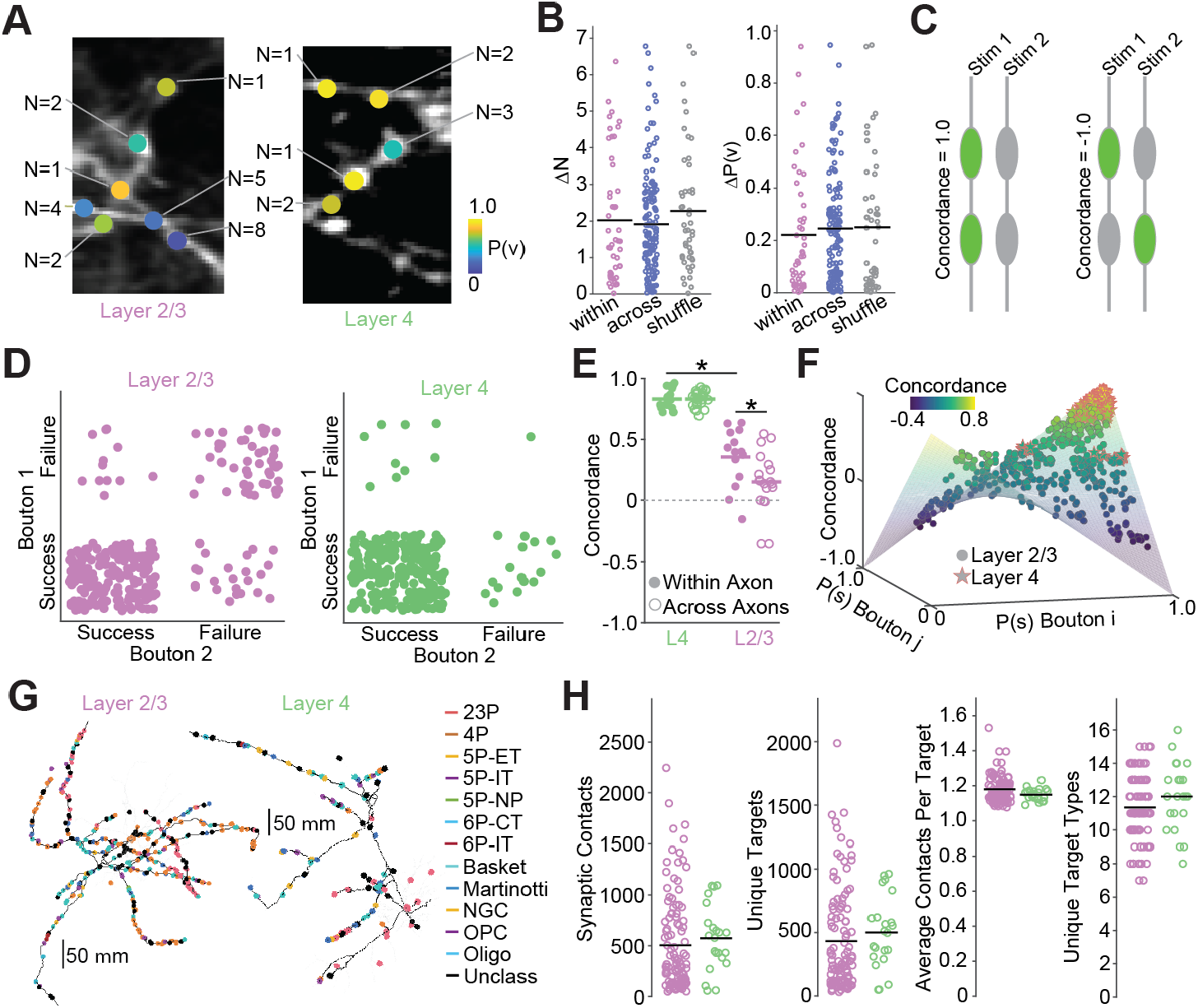
(A) Example of a layer II/III axon with multiple boutons and N(v) and P(v) overlaid on each ROI (left). The same for a layer IV axon (right). (B) Difference in N(v) between boutons on the same axon, across axons, and compared to shuffle (within axon mean 2.07±0.26 n=45 pairs, across axon mean 2.06±0.14 n=119 pairs) (left). Difference in P(v) between boutons on the same axon, across axons, and compared to shuffle (within axon mean 0.21±0.04 n=45 pairs, across axon mean 0.27±0.02, n=119 pairs). (C) Schematic of the concordance metrics (supplementary methods). (D) Example concordance quadrants for a representative layer II/III bouton pair (left, pink) and a layer IV bouton pair (right, green). (E) Concordance between pairs within an axon (closed circles; L23 within axon mean concordance 0.36±0.06) and between axons (open circles; L23 across axon mean concordance 0.15±0.06) for layer II/III pairs and layer IV pairs (L4 within axon mean concordance 0.83±0.02, L4 across axon mean concordance 0.84±0.01). Within axon concordance differed between L23 and L4 axons (p=3.48×10^-6^, two-sample ranksum test). (F) Hyperbolic paraboloid describing the total space of concordance with mean success rate across all pulses plotted against concordance for all bouton pairs. Layer II/III boutons are represented as circles and layer IV bouton pairs are represented as stars with a red outline. (G). Example axon skeletons from layer II/III (left) and layer IV (right) pyramidal neurons with boutons plotted as circles. Circle color indicates predicted post-synaptic cell type of a given connection. (H) From left to right: mean number of output boutons per cell for n=110 layer II/III and n=22 layer IV neurons classified as “interareal” proofreading status (L23 grand mean 505±45, L4 grand mean 574±68.6; p=0.135 two-sample ranksum test), number of unique post-synaptic partners per neuron (L23 grand mean 432±38.1, L4 grand mean 502±60.3; p=0.125 two-sample ranksum test), average number of boutons synapsing onto a post-synaptic partner per neuron (L23 grand mean 1.18±0.007, L4 grand mean 1.15±0.007; p=0.070 two-sample ranksum test), number of unique post-synaptic targets per neuron (L23 grand mean 14.7±3.4, L4 grand mean 12±0.43; p=0.176 two-sample ranksum test).

We next asked whether successful release events were correlated within single axons. For example, all same-axon boutons might succeed or fail to release for a given stimulus, consistent with some mechanism of shared influence. In contrast, each bouton might succeed or release independently, consistent with individual synapses acting fully independently from their neighbors.

We developed a concordance metric for each pair of boutons as ((#SS+#FF)-(#SF+#FS))/(Total #). The concordance value is 1 if each bouton in a pair succeeds or fails together, -1 if success and failure trials are perfectly inverted, and 0 if they are random. For our data, the average concordance was significantly greater than 0 for both populations with L2/3 boutons showing higher concordance within axon versus across (L2/3 within axon mean 0.36±0.06, across axon mean 0.15±0.06, p=0.021 two-sample ranksum test, L4 within axon mean 0.83±0.02, across axon mean 0.84±0.01, p=0.769 two-sample ranksum test). Notably, concordance rates were significantly higher for L4 versus L2/3 boutons on the same axon (p=3.5×10^-6^, two-sample ranksum test) (Figure 3E). For two independent synapses, concordance should be highest when each site has a very high or very low *Pv* (both release or fail for all APs, respectively). Indeed, our measured concordance values exhibited an expected parabolic relationship with the joint *Pv* values for each pair (Figure 3F). This result is further consistent with independent release across sites and highlights the variability with which a single neuron influences its different postsynaptic partners.

### Heterogeneity of postsynaptic targets may contribute to synaptic quantal variation

Synaptic release probability is known to vary for distinct connection types in the neocortex and hippocampus. For example, PN output to different interneuron classes differs in both synaptic strength and short-term synaptic plasticity (*33*–*35*). However, the overall diversity of synaptic targets for individual L4 or L2/3 PNs is unclear. We took advantage of the publicly available MICrONS serial electron microscopy volume ((*28*)) to quantify the distribution of postsynaptic partners receiving input from L4 or L2/3 PNs. For the proofread data set (110 layer 2/3 PNs and 22 layer 4 PNs), we found that, on average, cells in both layers form 500-600 total output synapses (layer 2/3 mean 504.76±44.04, layer 4 mean 574.27±68.60, p=0.135 two-sample ranksum test). Contrary to expectation, we observed that they also synapse onto a similar number of unique partners (layer 2/3 mean 432.33±38.15, layer 4 mean 501.77±60.29, p=0.125), giving an average of 1.18±0.007 synapses per post-synaptic target for layer 2/3 and 1.15±0.007 for layer 4 (p=0.069). Further, the average number of unique post-synaptic cell types was 11.35±0.19 for layer 2/3 neurons and 12.00±0.43 for layer 4 neurons (p=0.176). Thus, ultrastructural data corroborate functional heterogeneity and indicate that the output domain of pyramidal neurons displays an underappreciated complexity.

### Neuromodulatory control of synaptic release dynamics in vivo

The activity of cortical neurons is strongly influenced by variation in behavioral state (*36*–*40*). We therefore examined whether cortical synapses in vivo showed evidence of modulation over behaviorally relevant timescales. For all our experiments, we monitored the animal’s running speed on a wheel during imaging, dividing sessions into periods of quiescence or locomotion while simultaneously monitoring electrically evoked synaptic events from either L4 or L2/3 synapses (Figure 4A-B). We first quantified the relative amplitude of evoked events during locomotion versus quiescence, finding no change for L4 synapses (mean of 0.95±0.05 amplitude locomotion/amplitude quiescence, p=0.502 Wilcoxon Signed Rank test) but a significant reduction for those of L2/3 cells (mean of 0.85±0.02, p=1.3×10^-4^ Wilcoxon Signed Rank test)(Figure 4C). The reduced amplitude might arise from either smaller synaptic potency and/or increased number of failures. We therefore examined these metrics, finding that locomotion was similarly associated with a significantly reduced synaptic potency of L2/3 events (mean of 0.89±0.03, p=0.0027 Wilcoxon Signed Rank test) with no change in synaptic success rate (mean Δsuccess rate -0.04±0.02, p=0.058 Wilcoxon Signed Rank test)(Figure 4D). Thus, variation in behavioral state is associated with modulation of event-to-event glutamate concentration at single synapses.

Several prior studies indicate that behavioral state fluctuations are associated with fluctuations in acetylcholine (ACh) release (*36, 41*–*44*), and cholinergic receptors have also been shown to directly regulate glutamate release from cortical axons (*15, 42, 45, 46*). Thus, we investigated whether state-dependent modulation of release *in vivo* is shaped by cholinergic signaling. We imaged synaptic activity at L2/3 boutons following systemic injection of the muscarinic antagonist scopolamine, finding that drug administration blocked the state-dependent decrease in event amplitude and synaptic potency (mean of 0.97±0.04 amplitude locomotion/amplitude quiescence p=0.613 signed rank, mean of 1.00±0.04 potency locomotion/potency quiescence p=0.765 signed rank)(Figure 4C-D). Furthermore, scopolamine significantly increased the overall synaptic success rate relative to controls (0.83±0.02 scopolamine quiescent success rate versus 0.65±0.06 L2/3 no drug, p=0.014 ranksum test; 0.81±0.03 scopolamine locomotion success rate versus 0.61±0.05 layer 2/3 no drug, p=2.2×10^-4^ ranksum test) (Figure 4D). These data demonstrate that cortical synaptic transmission *in vivo* is under steady-state control of muscarinic signaling that strongly suppresses release probability.

**Figure 4.**
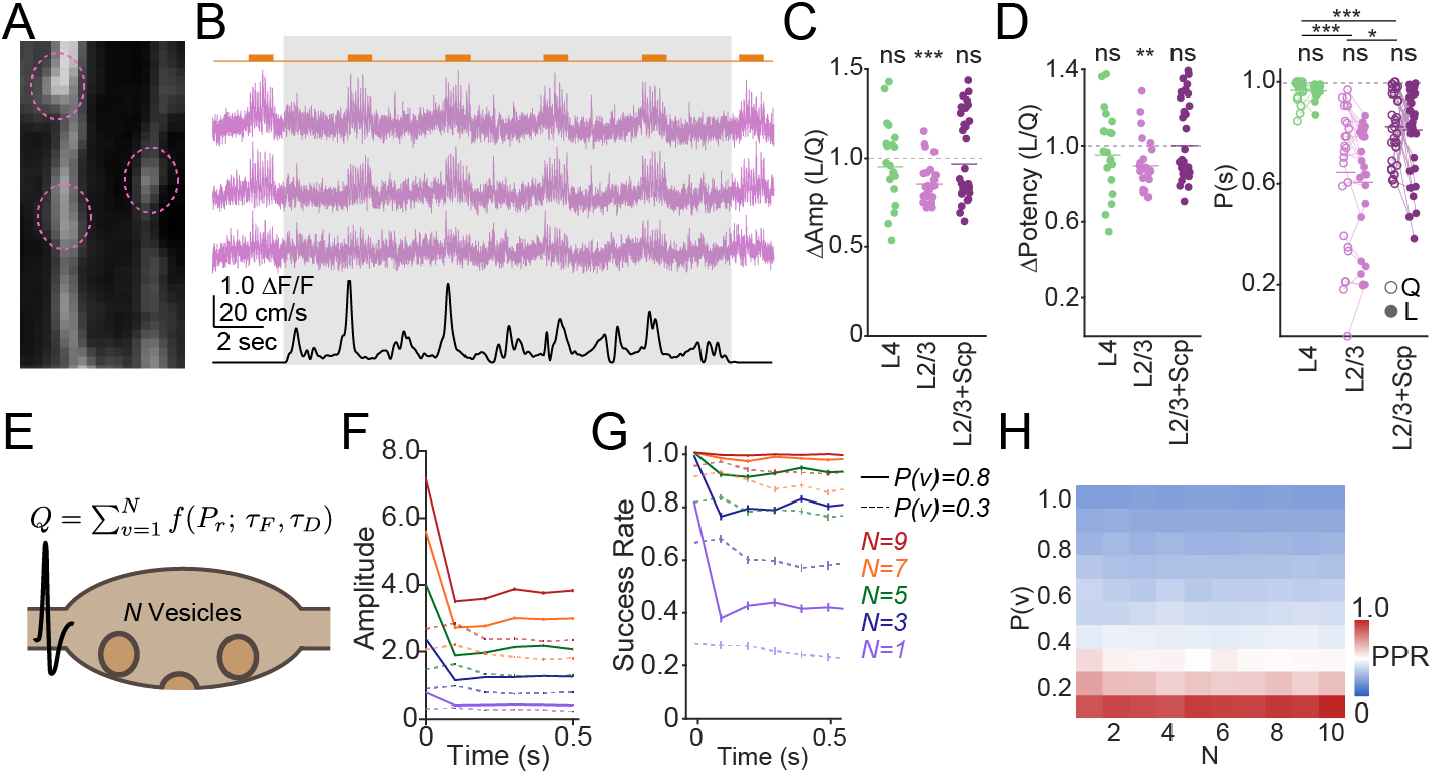
(A) Example of three layer II/III boutons imaged across quiescence and locomotion. (B) Traces from the three boutons in (A) with a grey box indicating a locomotion period. Wheel speed is plotted below. (C) Amplitude modulation (amplitude_locomotion_/amplitude _quiescence_) for layer IV boutons with no drug (mean 0.95±0.05, p=0.501 Wilcoxon Signed Rank test; n=20 boutons/2 animals), layer II/III boutons with no drug (mean 0.85±0.023, p=1.3×10^-4^ Wilcoxon Signed Rank test; n=25 boutons/3 animals), and layer II/III boutons with scopolamine (mean 0.97±0.04, p=0.614 Wilcoxon Signed Rank test; n=32 boutons/4 animals). (D) Left: potency by behavioral state for the same boutons. Layer 4 mean 0.95±0.05, p=0.502, layer 2/3 mean 0.89±0.03, p=0.0027, layer 2/3 with scopolamine mean 1.00±0.04, p=0.765. Right: success rate across for the same boutons. Layer II/III mean success rate locomotion: 0.65±0.06 and quiescence: 0.61±0.05; p=0.065 two-sample signed-rank test. Layer II/III with scopolamine mean success rate locomotion: 0.83±0.02 and quiescence: 0.81±0.03; p=0.985 two-sample signed-rank test. Layer IV mean success rate locomotion: 0.97±0.01 and quiescence: 0.97±0.01; p=0.296 two-sample signed-rank test. (E) Schematic illustrating short-term plasticity model applied to multivesicular release. Right: legend indicating N(v) and P(v) sweep in panels (H-I). (F) Amplitude for each of the N(v) and P(v) values in (G) right. (G) Mean success rate by pulse for the same N(v) and P(v) sweep. (H) Surface of a P(v) sweep demonstrating how normalized amplitude changes by initial release probability.

Our experimental data indicate that synaptic transmission at cortical layer 2/3 and layer 4 synapses is multi-quantal and with a range of vesicular release probabilities. We wondered whether our observations for cell type- and state-dependent synaptic dynamics emerged straightforwardly from these quantal metrics. To test this hypothesis, we constructed a model of synaptic release where *N* vesicles can be released independently, each with the same fusion probability *Pv*. The model is based on prior studies assuming independent time constants for synaptic facilitation and depression, arising from the decay of intracellular calcium concentration and kinetics of vesicle replenishing, respectively (*24, 47*) (Figure 4E). We explored the output of the model for a train of presynaptic impulses at 10 Hz, comparing a range of P(v) and N values. Results showed that changes in *Pv* exerted a strong influence over short-term synaptic plasticity, with higher release probability corresponding with greater short-term depression, while the number of vesicles had minimal impact on this metric (Figure 4F). In contrast, the number of vesicles strongly shaped the success rate for individual synapses, with higher number of vesicles supporting reliable transmission, even for extremely high-probability synapses (Figure 4G).

## Discussion

The quantal nature of synaptic transmission was initially established by recordings at the neuromuscular junction (*48, 49*), although several decades of research have confirmed this model at central synapses including those in the neocortex. However, controversy has remained over the number of vesicles able to fuse for each action potential, with studies providing evidence for both univesicular and multivesicular modes (*5, 6, 10, 50*– *52*). In most cases, multivesicular release was observed for synapses with high basal or experimentally elevated *Pv* (*8*–*10*), raising questions about its generalizability for connections between cortical PNs which have been reported to exhibit typical low *Pv* (*19, 53, 54*). Furthermore, all prior studies were carried out in ex vivo preparations, where ion concentrations, modulatory influences, and overall activity levels may not resemble conditions in the intact brain (*55*). Our data unequivocally demonstrate that MVR is observed in the intact, awake cortex. Moreover, we find that both L4 and L2/3 PNs exhibit MVR during spontaneous activity despite differing in their basal *Pv*.

The methodology we use in this paper completely isolates the pre-synaptic bouton and observes transmission at single synaptic contacts. These two aspects obviate the classical interpretational issues associated with recording whole-cell post-synaptic currents to infer pre-synaptic mechanisms. Namely, changes to post-synaptic receptor composition and averaging across multiple pre-synaptic contacts do not contribute to our measures of glutamate release. We anticipate that this strategy could be adopted in myriad basic and translational studies in which experimenters expect specific changes to the pre-synaptic compartment.

Functionally, MVR provides numerous computational advantages to synaptic transmission. First, it can greatly boost the amplitude of high *Pv* synapses, allowing several vesicles of glutamate to deliver a strong, high potency postsynaptic response. However, the presence of MVR also means that synaptic potency can vary as a function of *Pv*, greatly expanding the synaptic range of individual connections that can now transmit in a less binary, more analog fashion. Perhaps most importantly, MVR substantially increases the reliability of synapses, even under conditions of low *Pv* that might otherwise result in failure to release. Thus, synapses continue to deliver a signal, albeit weaker, rather than drop their transmission entirely.

Examining multiple boutons along the same axon further reveals that quantal parameters can differ greatly across boutons. This functional heterogeneity provides a scaffold over which individual neurons can tune their output at different synaptic contacts. This target-specific tuning has been widely implied by observations of differing short-term plasticity by post-synaptic cell type (*33*–*35, 56, 57*). Our work here provides a novel way that these phenomena could be instantiated at axonal boutons.

Our data also highlight a novel process by which cortical circuit dynamics can be regulated concomitantly with changes in behavioral state. Specifically, we find that muscarinic receptors exert a strong inhibitory control over glutamate release in vivo and that this modulation contributes to the state-dependent variation in synaptic strength. We show here that synaptic transmission itself is shaped by changes in arousal. Further, we observe that only a subset of boutons are responsive to the antagonist scopolamine, indicating that differential expression of muscarinic receptors may lend a dynamic heterogeneity to boutons that co-varies with behavioral state fluctuations. Overall, variation in presynaptic release properties including MVR may constitute a critical and underexplored mechanism for regulating circuit dynamics in vivo.

## Materials and Methods

### Animal Experiments

All animal experiments were approved by Institutional Animal Care and Use Committee of Yale University. Male and female adult mice were used in this study between post-natal day 60 (p60) and p200. Layer IV recordings were obtained from *Scnn1a*-Cre transgenic animals (JAX Strain #009613) and layer II/III recordings were obtained from wild-type C57BL/6J animals. Transgenic and wild-type animals were bred within the Yale Animal Research and Care facility. Mice were housed in 12h:12h light/dark cycles in a climate-controlled vivarium throughout all experiments. Experiments were performed during the light period. No statistical tests were conducted to determine sample sizes prior to the study.

### Surgical Procedures

Two-photon headplate and cranial window surgery was performed as previously described (*58, 59*). Briefly, mice were anesthetized using 5% isoflurane for induction and 2% isoflurane for maintenance throughout the procedure. A custom titanium headplate was attached to the rostral skull between the eyes and secured with CB Metabond. For layer IV bouton imaging, 800 nanoliters of AAV9-CAG-FLEX-iGluSnFR4f-PDGFR (Addgene #234450) at 5×10^12^ GC/mL was injected at 350 micrometers below the dural surface. For layer II/III imaging, AAV1-CaMKII-Cre (Addgene #105558) at 5×10^9^ GC/mL was co-injected alongside the iGluSnFR-containing virus at 200 micrometers below the dura. A 3mm by 3mm craniotomy was then drilled over the injection site and the skull was removed with the dura kept intact. A small incision was made in the dura away from the injection site and a bipolar stimulating electrode (P1 technologies MS303/2-B/SPC) was lowered into the brain at a 15-20 degree angle with the tip reaching approximately the injection location. A double-layered glass coverslip was then affixed over the brain and electrode using cyanoacrylate glue and the window and remaining skull surface were sealed with Metabond. Animals were allowed to recover for at least 3 weeks before imaging began.

### Two-Photon Imaging

Animals were acclimated to head fixation prior to the start of imaging. Awake mice sat and ran freely on an unmotorized wheel throughout each imaging session. Two-photon imaging was conducted using a custom-built resonant-galvo-scanning microscope fed by a SpectraPhysics MaiTai femtosecond pulsed laser tuned to 920nm. All recordings were conducted between 250-280Hz frame rate. Single-bouton imaging was conducted at 15-30 optical zoom. ScanImage software was used to control microscope hardware components. Behavioral variables were collected via a magnetic wheel sensor and infrared videography and digitized using Cambridge CED acquisition hardware with Spike2 software.

### Electrical Stimulation

Electrical pulses were delivered to the cortex via the implanted bipolar stimulating electrode connected to an ISO-Flex stimulus isolator. The amplitude was tuned in pilot experiments to the minimum necessary to reliably evoke calcium activity in soma and axons reported by jGCaMP8f. 100 microsecond duration pulses were delivered in groups of 10 pulses at 10Hz to assay Pr and STP. A recovery pulse followed the initial train by 1 second. No behavioral signs of pain or stimulus perception were observed in any of the animals.

### Pharmacological Experiments

For experiments involving *in vivo* pharmacology, scopolamine hydrobromide was dissolved to a final concentration of 1 mg/mL in saline. Animals were weighed before injection and the injection volume was tuned to deliver 1 mg/kg intraperitoneally. Glutamate recording was started 10-15 minutes after initial injection. No signs of pain or distress were observed after injection. Successful administration of scopolamine was confirmed by verifying that the pupil dilated at room light levels.

### Data Analysis

Raw movies were motion corrected using NormCorre (*60*). Recordings typically contained 1-10 boutons per movie. ROIs were manually selected and shapes were refined and separated using seeded constrained non-negative matrix factorization (CNMF) implemented through the CaImAn two-photon analysis package (*61*). Traces were extracted as the mean fluorescent value of all pixels within an ROI per frame. All subsequent analysis of two-photon imaging data and behavioral variables was conducted using custom-written functions in MATLAB.

### Event Detection

Events were detected from raw glutamate traces in a multi-step process beginning with minimizing the sum of squared errors between a template waveform and a sliding window of the trace ((*62*)). This process was repeated once more after refining the template based on the average of the first round of detection. Each detected event was extracted in a window of -20ms to +50ms around the detected timestamp and then subjected to single-exponential fitting. Candidate waveforms that fit an exponential decay from the peak onwards with r^2^ > 0.25 were included in the analysis. All events were manually inspected and false positives/negatives were corrected. Amplitudes of spontaneous events were calculated as ΔF/F_0_ using the sample immediately before the rise onset as the F_0_ value. In order to avoid overestimating amplitudes due to accumulation on previous transient decays, amplitudes of evoked events were calculated as the difference between the peak sample and an extrapolated single-exponential decay fit from the peak of the previous stimulated transient; the first stimulated transient used subtraction from the 200ms baseline before the train onset. Event frequency was calculated as the number of events occurring over the entire recording divided by the total recording duration.

### Multiquantal Determination

To classify boutons as multi- or uniquantal from spontaneous data, event amplitudes were normalized to the minimal quantum (“q”) estimated as the mean of a gaussian reflecting the left tail of the amplitude histogram. The mean of the left-tailed gaussian fit was taken as q.

Histograms of normalized amplitudes were then fit with either a single gaussian or iteratively with up to 5 gaussians centered on multiples of q. Likelihood Ratio Test (LRT) and Bayesian Information Criterion (BIC) were then calculated based on the model fit to determine whether a single or multiple gaussians fit significantly better. Boutons in which multiple gaussians were determined to fit better via both LRT (p < 0.05) and BIC (ΔBIC > 2) were classified as multiquantal. As an independent verification, the noise standard deviation was calculated from event-free periods throughout the trace and overlaid on the amplitude histograms at 1q. Boutons classified as “uniquantal” displayed amplitude distributions entirely within this 1q + noise standard deviation overlay; boutons classified as “multiquantal” had right-tailed amplitude peaks outside of the noise estimate.

### Behavioral State Partitioning

Locomotion was recorded simultaneous to two-photon imaging using a rotational magnetic sensor affixed to the wheel. The magnetic sensor signal was aligned to the imaging frames via the galvo triggers recorded simultaneously on Spike2. The raw wheel signal was then unwrapped and converted to units of cm/s. Speed data were thresholded at 5 cm/s to identify locomotion onsets and offsets. The wheel signal was then binarized as locomotion on/off. Locomotion or quiescence bouts lasting less than 2 seconds were excluded from the analysis. For all remaining bouts, glutamate events were classified as either occurring during locomotion or during quiescence for behavioral state-based analyses.

### Quantification of Electrically Evoked Activity

Event detection was conducted independently of stimulation times. After event detection, events were assigned as belonging to a given stimulation if they occurred within 50 milliseconds (∼12 samples) of a stimulation time. Failures were quantified as the proportion of stimulation times which did not have an associated glutamate event. As an independent verification, raw glutamate traces from a subset of recordings were analyzed around stimulation times and failures were classified as the peak of the post-stimulus window failing to exceed 3 standard deviations of the baseline standard deviation from 0 to -200 milliseconds before stimulus onset.

Short-term plasticity was defined as the ratio of the event amplitude of pulse 2-11 and pulse 1 for a 10Hz stimulus train. For analysis of potency, failure trials were excluded from the average.

### Estimation of Number of Release Sites and Per-Site Release Probability

Electrically evoked recordings were used to estimate N and P(v) using two independent methods.

In one method, we adapted a binomial model used in (*30*) that estimates the number of release sites as a function of trial-to-trial variability in single stimulation pulses (equation 1). As N is expected to be a structural variable invariant to STP, we separately calculated N for each stimulation pulse in the 10Hz and took the mean N value obtained across pulses as the N for a given bouton.

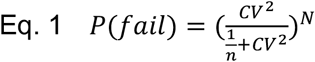

The other method involved examining the difference between success rates and normalized amplitude across pulse pairs within the train. The ratio of the success rates should equal the ratio of the normalized amplitudes at N=1. If N is greater than 1, the depression observed in the amplitudes will be greater than the depression in the success rate across pulses. Values of N can then be swept to determine which value results in correspondence between the two ratios (equations 2-4). As this relationship holds regardless of the sequence of pulses, we conducted the analysis across all possible pairs and determined N for a given bouton as the mean value obtained across all pairs. The N sweep was conducted from 0.1 to 20 potential N values in steps of 0.25. Values pinned to the floor or ceiling of the N search grid (both chosen as sub- or supraphysiological values) were excluded from analysis and the N for that bouton was left as undetermined by the method. For both methods, N values were rounded to the nearest integer before back-calculating P(v).

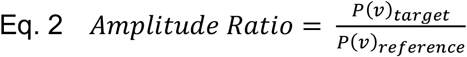

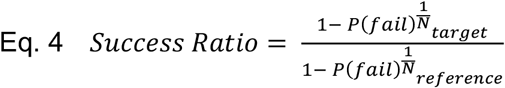

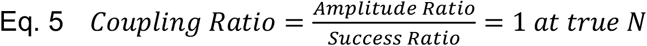

After obtaining N for a given bouton, we back-calculated P(v) for each pulse across the train per bouton as 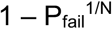 assuming a uniform release probability across all sites.

### Segmentation of Axons from Imaging Fields of View

To determine which boutons in a given field of view are from the same axon versus different, we employed an entirely structural approach. We opted for this to avoid circularity in segmenting based on activity and then later examining activity correlations. In addition, we observed decorrelated activity from boutons that were visually on the same axon.

Movies were collapsed down to their average projection across frames and filtered using a Frangi filter of span [0 1] to accentuate tubular structures. The filtered image was then thresholded to create a binary mask and skeletonized. Skeletons were pruned to eliminate artifactual branches with less than 5 pixel length. ROIs were assigned as belonging to a particular axon if the ROI centroid was no more than 3 pixels away from any point on a given axon skeleton. All segmentation was manually corrected.

### Concordance Between Boutons

Concordance was defined according to equation 6 for 275 total stimulation pulses across 25 10Hz stimulus trains. Electrically evoked activity was analyzed between bouton pairs determined to share an axon and across axons within a field of view. For each trial, successes and failures were mapped in quadrant grids exemplified in figure 3C-D. The total concordance value for a bouton pair was then calculated from the total successes and failures in the grid.

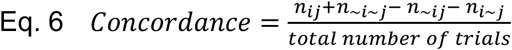

To visualize how bouton pairs fill the possible space of concordance, we plotted a hyperbolic paraboloid (equation 7) in three dimensions and overlaid the mean success rates and calculated concordance for all bouton pairs within and across individual axons.

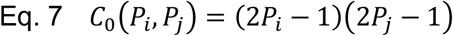

### Analysis of the MICrONS Volume Electron Microscopy Dataset

The neuron ID of all layer II/III pyramids was obtained by querying the Minnie65 dataset through the Python API. Of all the neurons classified as “L23P”, we selected only those with a proofreading status of “interareal” as of November 2025 to only analyze reconstructions that were as complete as possible and, critically, to ensure that we always analyzed neurons with an unambiguous axon path between the soma and any distal branch. We then obtained all of the unique synapse ID for any synapse where a proofread layer II/III ID was listed as “pre-synaptic” to index each unique neuron in our analysis (110 layer II/III pyramids) with their boutons (∼55000 unique boutons). Each synapse was associated with a pre-synaptic partner ID (i.e. being a bouton on this cell), a post-synaptic partner ID and putative cell type, and XYZ coordinates within the volume. We also downloaded the available neuron mesh for each neuron to be used for axonal reconstruction. Subsequent analyses were conducted in MATLAB using custom written functions.

We used a combination of the bouton XYZ coordinates and the three-dimensional neuron mesh to reconstruct the axon skeleton. From each bouton location, we expanded the skeleton out by taking the centroid of the mesh section perpendicular to the mesh surface (the least ellipsoid slice). We also estimated the soma position as the largest ellipsoid volume of the mesh. Skeletonization and soma determination were visually verified in a subset of neurons.

From the skeleton, branching was identified as any segment of the skeleton with more than converging skeleton segments. For post-synaptic cell type interspersion analysis, each branch was analyzed independently. Each branch received a polarity based on proximity to the bifurcation point defined as being the soma side. Interspersion was calculated as the total number of transitions between any given pair of post-synaptic cell types divided by the total number of transitions on a branch. Transition probabilities were then pooled across branches and then across neurons. Enrichment over chance was calculated as the proportion of a given transition divided by the number of such transitions occurring in 5000 shuffles per branch.

### Modeling Short-Term Dynamics and Release Probability

We adapted the Abbott & Regehr short-term plasticity model (*24, 47*) of release to determine how multiquantal release shapes synaptic output. As per the standard model, we calculated the per-trial release probability for a given impulse as a function of facilitation and depression on basal P_release_ (equations 8 and 9).

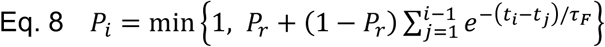

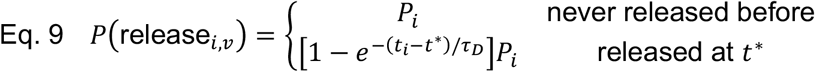

We then adapted the model to calculate an independent P(v) per trial for each of N independent release sites (equation 10). Time constants for facilitation and depression were set to 0.05 and 0.25 seconds, respectively, to match the STP curves obtained from our evoked glutamate recordings. We then ran the model for 1000 iterations of 6 spikes with 100ms inter-spike interval, mimicking our 10Hz *in vivo* stimulation.

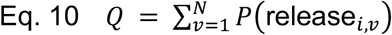

